# Leakage-Controlled Benchmarking of Variant Effect Predictors and Development of Gradient-Boosting Consensus Model

**DOI:** 10.64898/2026.10.30.763022

**Authors:** Martin Havlasek, Petr Kabourek, Milos Musil, Jiri Damborsky, Stanislav Mazurenko, David Bednar

## Abstract

Single amino acid mutations are among the most common forms of genetic variation, with effects ranging from negligible to severe disruption of protein function. Accurate quantitative evaluation of protein variants therefore represents a key challenge in medicine and protein engineering. Computational variant effect predictors (VEPs) are widely used to prioritize missense variants, yet their fair comparison is hindered by data circularity, benchmarks restricted to human clinical annotations, and limited availability of code and training data, all of which can inflate apparent performance. Here, we reviewed 18 recent and 3 older, widely adopted variant effect predictors for reproducible benchmarking and assembled a rigorously curated, leakage-controlled *Benchmark dataset* by integrating clinically annotated human variants from ClinVar and Humsavar as well as cross-species variants with functional annotations. The evaluation on this dataset revealed a narrow overall performance range for the benchmarked tools (AUC 0.631–0.758) and a consistent drop from clinical to functional data. Surprisingly, several tools published two decades ago demonstrated comparable performance to mid-ranked modern predictors. Building on these results, we developed a gradient-boosting consensus model, PredictSNP^prot^, which achieved an AUC of 0.82 on a held-out test set, outperforming the original PredictSNP meta-predictor and the best individual state-of-the-art tool by 13.6 and 5.7 percentage points, respectively. The *Benchmark dataset*, including the predictions from the evaluated tools, is available at https://doi.org/10.5281/zenodo.23597130 and PredictSNP^prot^ consensus predictor is available at https://github.com/loschmidt/PredictSNPprot.

**Key Points:**

- We constructed a leakage-controlled benchmark dataset of 10,385 clinical and cross-species functional variants, thoroughly filtered to exclude overlap with training data of benchmarked tools.
- The benchmarking of 14 sequence-based variant effect predictors on this dataset yielded narrow AUC range (0.631–0.758) and most performed markedly worse on functional than on clinical variants.
- We developed PredictSNP^prot^, a gradient boosting consensus model, integrating 9 constituent tools and reaching an AUC of 0.818 on a held-out test set, compared with 0.761 for the best single tool.
- The benchmark dataset including all in-house generated tool predictions as well as the PredictSNP^prot^ code are openly available at Zenodo and GitHub, respectively.

## Introduction

Mutations are fundamental drivers of evolution, responsible for sequence diversity that enables organisms to adapt to changing environmental conditions^1^. In coding regions, much of this variation comes from single nucleotide polymorphisms (SNPs), which may lead to amino acid substitutions in the encoded proteins^2^. The effects of these missense variants span a broad spectrum, ranging from well-tolerable neutral or beneficial changes to deleterious alterations that disrupt protein structure, stability, or function.^3,4^ Understanding these effects is therefore essential for identifying pathogenic variants in human disease^5^ and for evaluating substitutions in protein design^6^.

Methodological advances and the dramatic cost reduction of sequencing technologies have enabled the generation of massive sequence datasets^7–11^, paving the path to the identification of disease-causing mutations in patients with genetic disorders and highlighting variants with significant impact on protein fitness. However, experimental evaluation of the functional effects of these variants remains challenging^12^. Despite progress in high-throughput experimental assays such as multiplexed assays of variant effect^13^ (MAVE) and deep mutational scanning^14^ (DMS), these approaches remain difficult to scale to the proteome level, and their accuracy and consistency in capturing functional phenotypes are still limited^12,15^, leaving most variants classified as being of uncertain significance^16^.

Over the past decades, numerous computational variant effect predictors (VEPs) have been developed to help annotate the vast number of experimentally unclassified protein variants. Although these methods have not yet achieved the accuracy required for decisive classification in clinical settings under American College of Medical Genetics and Genomics and the Association for Molecular Pathology (ACMG/AMP) guidelines^17^, they have earned their place as valuable prioritization tools and supplementary classification evidence, offering a rapid and scalable alternative to costly experimental assays. VEPs typically exploit various biological, chemical, physical, and evolutionary properties derived from protein sequence or structure to estimate the likelihood of a given mutation being deleterious. Structural information has been shown to provide valuable context for mutational impact prediction^18^, as demonstrated by PrimateAI-3D^19^, which directly uses structural input, and AlphaMissense^20^, which learns structural context through an AlphaFold2^21^-derived architecture without using a structure as input. Nevertheless, the limited availability of high-quality structures relative to sequences and the convenience of sequence-only input have led the vast majority of VEPs to be sequence-based.

Traditional sequence-based VEPs can be broadly categorized into three groups: (i) evolutionary-based methods, such as GEMME^22^, PHACT^23^, and PROVEAN^24^, which rely primarily on amino acid conservation derived from multiple sequence alignments; (ii) machine learning-based methods, employing either supervised (e.g., 3Cnet^25^, MutPred2^26^, PolyPhen-2^27^) or unsupervised (e.g., EVE^28^) approaches to extract mutation-specific features; and (iii) methods based on protein language models (pLMs, e.g., ESM1b^29^), which apply techniques from natural language processing to model the likelihood of all possible sequence variants. More recent VEPs often combine these strategies; for example, VariPred^30^, VESPA^31^, and VespaG^32^ integrate pLM embeddings with supervised learning, while PHACTboost^33^ combines evolutionary conservation with gradient boosting. Finally, many methods aggregate outputs from multiple independent VEPs within consensus-based workflows (i.e., meta-predictors), an approach that has been shown to generally outperform individual predictors and yield more reliable results^34,35^. We have previously developed the PredictSNP^36^ webserver, which integrates eight individual prediction tools, six of which contribute to its consensus score. Despite its wide adoption by the community, demonstrated by processing over 80,000 jobs and collecting over 800 citations since its release in 2014, the substantial number of newly developed, advanced VEPs present an opportunity for significant improvements in its predictive accuracy.

Despite significant improvements in newly developed VEPs demonstrated by various benchmarks^35,37,38^, their direct, comprehensive, and unbiased comparison remains challenging. Two types of data circularity are the most prominent causes of this issue, leading to overly optimistic estimates of tool performance^38,39^. Type 1 circularity arises directly from data leakage between the training datasets of supervised methods and the testing datasets used for benchmarking^37^. Type 2 circularity stems from features derived from specific sequence regions, proteins, or protein families, indirectly biasing predictions toward skewed outcomes when the method encounters similar targets during inference^37,39^. Preventing these issues is challenging, since training data are often unavailable and clinically labeled data used for benchmarking are limited to a few strongly overlapping sources, such as ClinVar^40,41^, UniProt’s^42^ Humsavar, and HGMD^43^. Moreover, most benchmarks rely on precomputed predictor scores deposited in databases such as dbNSFP^44,45^, limiting control over the evaluation process.

In this study, we therefore aimed for fair, unbiased benchmarking by addressing these limitations directly. First, from the total set of 18 reviewed, recently published predictors, we identified 11 VEPs that meet strict reproducibility and transparency criteria, enabling local installation and computation of the predictions in-house rather than relying on precomputed scores. We further extended the list of selected tools by including 3 older and widely adopted VEPs, yielding a final set of 14 tools for the benchmark. Second, we constructed a comprehensive *Benchmark dataset* combining human and non-human variants from clinical annotations and functional deep mutational scanning (DMS) assays, thoroughly filtered to exclude any overlap with predictors’ training data at the level of specific variants while monitoring the overlap at the level of full-length protein sequences. Using this strategy, we conducted thorough benchmarking of the selected VEPs, spanning diverse methodological foundations, including purely evolutionary-based methods, pLM-based tools, and supervised predictors. Finally, by combining this validated set of VEPs, we developed PredictSNP^prot^, a new consensus model based on gradient boosting that yields robust protein missense effect predictions surpassing the accuracy of the original PredictSNP meta-predictor and all individually tested variant effect predictors.

## Materials and methods

### Selection of the Variant Effect Predictors

Protein sequence-based variant effect predictors (VEPs) released between 2015-2024 were identified through a thorough literature review, and those satisfying the following criteria were selected for evaluation: (i) availability of a stand-alone, installable version; (ii) support for user-defined protein sequences as input (by default or upon minor source code modifications); and (iii) availability of training data (for supervised methods). Predictors requiring gene or chromosomal identifiers, rather than user-defined protein sequences, as input were excluded from consideration. Some otherwise eligible tools were subsequently excluded due to limitations hindering installation or calculation on the *Benchmark dataset*, such as a requirement for manual computation of input features or excessive computational demands. Additionally, the original PredictSNP^36^, PROVEAN^24^, and PolyPhen-2^27^ were included as baselines despite being released prior to 2015. Full selection criteria and the complete list of tools considered, including those excluded and the corresponding reasons, are provided in **Table S1**.

### Benchmark dataset construction

#### Raw dataset

The *Raw dataset* was composed from four distinct data sources: ClinVar^40,41^, UniProt’s^42^ Humsavar, Protein Mutant Database^46^ (PMD), and massively mutated proteins^36,47–49^ (MMP). Only records providing a specific protein identifier (UniProt or GenBank ID or equivalent), a defined amino acid substitution, and an experimental annotation clearly distinguishing between neutral and disruptive effects were considered. ClinVar data (variant_summary.txt, version July 2024) were downloaded from the NCBI ClinVar FTP repository. Only records annotated as “single nucleotide variant” in the Type field were retained. Gene identifiers (NCBI RefSeq accessions) and mutation descriptions were extracted from the Name field using the HGVS protein-level notation. ClinVar variants were binarized using the ClinicalSignificance field, with the ACMG/AMP germline classification as the decisive criterion. Entries whose classification was pathogenic or likely pathogenic were assigned label 1, and entries classified as benign or likely benign were assigned label 0. Composite entries containing multiple ACMG/AMP terms (e.g., “Pathogenic/Likely pathogenic”) or additional annotations (e.g., risk factor, drug response, association, low penetrance) were assigned according to their ACMG/AMP classification, and the additional annotations did not affect the assignment. Two entries without an ACMG/AMP classification were assigned explicitly: “Likely risk allele” was assigned label 1, and “protective” was assigned label 0. Variants of uncertain significance and those with conflicting classifications were excluded. Finally, mutations corresponding to termination codons or nonspecific substitutions (i.e., those beginning or ending with ‘X’ or ‘–’, reflecting “Ter” or “Xaa” residues) were filtered out to retain only specific missense substitutions.

For Humsavar, humsavar.txt (version 29-May-2024) was downloaded from the UniProt FTP repository. UniProt accessions served as protein identifiers, and corresponding sequences were retrieved from UniProt. Variants labeled “LB/B” (likely benign/benign) were assigned class 0, those labeled “LP/P” (likely pathogenic/pathogenic) were assigned class 1, and entries labeled “US” (uncertain significance) or otherwise unmapped were excluded. For both ClinVar and Humsavar, corresponding full-length protein sequences were subsequently retrieved from GenBank/RefSeq and UniProt, respectively, using the standardized identifiers extracted above. Data from PMD and MMP required only minor formatting adjustments to ensure consistency with the other sources, as these datasets had already been curated, sequence-mapped, and preprocessed during development of the original PredictSNP^36^.

Entries from all four sources were merged into a unified dataset retaining source annotations. The dataset was sanitized to retain only complete entries with valid protein sequences (i.e., excluding those with sequences containing wildcard symbols or other ambiguities) and with correct alignment between wild-type amino acid from the mutation and the corresponding residue at the annotated sequence position, yielding the *Raw dataset* (257,112 records).

#### Filtered dataset

The *Raw dataset* was then filtered in two steps. First, duplicate entries and records with conflicting labels (i.e., identical mutations assigned different labels across sources) were removed. Second, to prevent overlap with the training data of the evaluated VEPs, we collected the training datasets of all tested supervised machine learning-based predictors considered: PredictSNP, 3Cnet, Envision, MutPred2 (publicly available data only; HGMD-derived training data were unavailable), PHACTboost, PolyPhen-2, VariPred, VESPA, VESPAl, and VespaG. Each dataset was reformatted into a standardized scheme (identifier, mutation, sequence) using the retrieval approach described above, giving 4,949,968 training records across 43,401 unique protein sequences. The *Raw dataset* was then cross-referenced against these records, and any entry corresponding to a specific variant (i.e., a specific mutation of a specific sequence) present in any tool’s training data was removed. Where a training record named no specific variant (the VespaG training data contain only protein sequences), all variants of that sequence were removed. Each remaining entry was then assigned to the strictest independence tier it satisfied: (i) *clean sequence* (12,452 records), where none of the protein’s mutations appeared in training data; (ii) *clean site* (65,192 records), where the protein was present in training data but the specific mutation site was not; or (iii) *clean variant* (19,504 records), where both the protein and the mutation site occurred in training data, but not the specific substitution. Tiers are therefore mutually exclusive, although a *clean sequence* record also meets the criteria of the two less strict tiers. This procedure yielded the *Filtered dataset* (97,148 records; 10,549 unique sequences).

#### Benchmark dataset

The *Filtered dataset* was subsequently rebalanced to achieve comparable representation across substitution types and pathogenicity classes. Balancing was performed independently for each data source to ensure comparable subset sizes in the final dataset. Selection followed a hierarchical strategy, prioritizing entries from the *clean sequence* subset, followed by progressive enrichment with *clean site* and *clean variant* entries, up to a maximum of ten entries per unique substitution-label combination per source, with unique protein sequences prioritized to maximize diversity. This procedure yielded the final *Benchmark dataset* (10,385 records), used for evaluation of the individual VEPs. Detailed composition of all three datasets is provided in **Table S2**. *Raw, Filtered*, and *Benchmark datasets* are available at https://doi.org/10.5281/zenodo.23597130.

### Implementation and benchmarking of VEPs

The selected tools were installed locally as Singularity image containers and configured to accept user-defined protein sequences as input. Default parameters were used unless otherwise stated. Full configuration details, including models, parameters, and code modifications, are provided in **Table S3** and sequence databases used are specified in **Table S4**. Multiple sequence alignments (MSAs) for GEMME^22^, PHACT^23^, and PHACTboost^33^ were generated using an externalized workflow derived from the PHACT pipeline (see **Supplementary Note 1** for details). Predictions for the *Benchmark dataset* were computed using the MetaCentrum Czech National Grid Infrastructure, except for PredictSNP, which was accessed via its webserver API. Upon completion, predicted scores for all tools were retrieved and compiled for downstream evaluation.

The area under the receiver operating characteristic curve (AUC) was used as the primary performance metric. Additional confusion-matrix-derived metrics, including accuracy, precision, recall, F1 score, and Matthews correlation coefficient (MCC), were calculated using a tool-specific classification threshold derived from Youden’s J statistic. All metrics were computed using the scikit-learn Python package (v1.8.0)^50^ and applied to the *Benchmark dataset*’s common subset of variants for which all tested tools produced a prediction. In addition, performance was evaluated separately on subsets stratified by data source and by level of independence from the tools’ training data (*clean sequence, clean site*, and *clean variant*), allowing assessment of performance degradation under stricter independence criteria. Additionally, the performance was evaluated individually for each tool on the subset of *Benchmark dataset*, where the tool produced a valid prediction. The failure rate, defined as the fraction of benchmark entries for which a tool failed to produce a prediction, was recorded for each tool as an additional measure of practical usability and robustness.

### Consensus model development and validation

#### Selection of integrated VEPs

Thirteen benchmarked tools (**Table 1**) were considered for integration within the consensus model (PredictSNP was excluded directly as it was superseded by the new consensus model). To ensure practical usability of the meta-predictor, we excluded VEPs, which exhibited an observable high fraction of entries for which they failed to produce a prediction on the *Benchmark dataset* (observed failure rate > 2%, EVE, PolyPhen-2; and VESPA see **Table 2** for details). Additionally, MutPred2 was excluded from the pool of constituent tools due to inability to perform saturation mutagenesis predictions (i.e., predictions for all possible single-point substitutions) in reasonable runtime while the included VEPs produced saturation mutagenesis by default or their runtime scaled efficiently with exhaustive list of all possible mutations (PROVEAN, 3Cnet, and VariPred), rendering such computations feasible.

**Table 1:** Overview of variant effect predictors included in the benchmark. The table provides the publication year, general categorization, underlying computational principle, and primary reference. Tools are grouped into recently developed predictors (published 2015–2024) and legacy predictors (published before 2015), retained as community-standard baseline comparators^*^.

| Tool | Published | Category | Principle | Reference |
|---|---|---|---|---|
| **GEMME** | 2019 | Evolutionary/MSA | Unsupervised epistatic modeling of evolutionary fit and frequency from MSA | [<sup>22</sup>] |
| **MutPred2** | 2020 | Evolutionary/MSA + Supervised machine learning | Bagged ensemble of feedforward neural networks on structural, functional, and evolutionary sequence features | [<sup>26</sup>] |
| **3Cnet** | 2021 | Supervised machine learning | Multi-task RNN on sequence context, jointly trained on clinical variants and MSA-derived synthetic conservation data | [<sup>25</sup>] |
| **EVE** | 2021 | Evolutionary/MSA | Bayesian variational autoencoder inferring an evolutionary index from MSA | [<sup>28</sup>] |
| **PHACT** | 2022 | Evolutionary/MSA | Phylogeny-aware tolerance scoring from ancestral reconstruction probabilities | [<sup>23</sup>] |
| **VESPAl** | 2022 | pLMs + Supervised machine learning | Logistic regression on ProtT5-derived conservation predictions alone (lightweight VESPA variant) | [<sup>31</sup>] |
| **VESPA** | 2022 | pLMs + Supervised machine learning | Logistic regression ensemble on ProtT5-derived conservation predictions and substitution log-odds ratios | [<sup>31</sup>] |
| **ESM1b** | 2023 | pLMs | Zero-shot log-likelihood ratio from a masked protein language model | [<sup>29</sup>] |
| **VespaG** | 2024 | pLMs + Supervised machine learning | Feedforward network on ESM-2 embeddings, trained on GEMME-derived pseudo-labels | [<sup>32</sup>] |
| **VariPred** | 2024 | pLMs + Supervised machine learning | Feedforward neural network on ESM-1b protein language model embeddings | [<sup>30</sup>] |
| **PHACTboost** | 2024 | Evolutionary/MSA + Supervised machine learning | Gradient boosting on PHACT scores combined with MSA and phylogenetic features | [<sup>33</sup>] |
| **PolyPhen-2** | 2010 | Supervised machine learning | Naive Bayes classifier on sequence and structural features | [<sup>27</sup>] |
| **PROVEAN** | 2012 | Evolutionary/MSA | Delta alignment score from similarity-weighted clustering of homologous sequences | [<sup>24</sup>] |
| **PredictSNP** | 2014 | Consensus | Consensus meta-predictor integrating six individual tools | [<sup>36</sup>] |

<sup>\*</sup>Full selection criteria and the complete list of tools considered, including those excluded and the corresponding reasons, are provided in Table S1. The source of code, its modifications, and configurations are detailed in **Table S3**. Abbreviations: pLMs - protein language models; RNN – recurrent neural network, MSA -. multiple sequence alignment

**Table 2:** Overall performance of the 14 benchmarked VEPs on the Benchmark dataset. Calculated is the percentage of the 10,385 Benchmark entries for which a tool returned a prediction. All performance metrics were computed on the common subset, i.e., the entries for which all 14 tools returned a prediction (N = 8,622). The area under the receiver operating characteristic curve (AUC) was the primary metric. Accuracy, precision, recall, F1 score, and Matthews correlation coefficient (MCC) used a classification threshold derived from Youden’s J statistic on the common subset. Tools are ranked by AUC.

| **Tool** | **Calculated (%)** | **AUC** | **Accuracy** | **Precision** | **Recall** | **F1** | **MCC** |
|---|---|---|---|---|---|---|---|
| VariPred | 99.99 | 0.758 | 0.692 | 0.767 | 0.649 | 0.703 | 0.393 |
| VESPA | 97.56 | 0.755 | 0.693 | 0.742 | 0.697 | 0.719 | 0.382 |
| PHACT | 98.34 | 0.746 | 0.695 | 0.761 | 0.669 | 0.712 | 0.395 |
| GEMME | 98.31 | 0.737 | 0.688 | 0.719 | 0.733 | 0.726 | 0.365 |
| VESPAl | 99.45 | 0.734 | 0.682 | 0.723 | 0.705 | 0.714 | 0.357 |
| ESM1b | 99.99 | 0.732 | 0.680 | 0.761 | 0.630 | 0.689 | 0.373 |
| VespaG | 98.78 | 0.731 | 0.672 | 0.739 | 0.647 | 0.690 | 0.349 |
| MutPred2 | 99.99 | 0.729 | 0.678 | 0.727 | 0.687 | 0.706 | 0.352 |
| PolyPhen-2 | 95.65 | 0.722 | 0.671 | 0.728 | 0.664 | 0.694 | 0.342 |
| PHACTboost | 98.40 | 0.718 | 0.679 | 0.702 | 0.749 | 0.725 | 0.343 |
| PROVEAN | 100.00 | 0.708 | 0.681 | 0.686 | 0.802 | 0.739 | 0.343 |
| EVE | 86.04 | 0.700 | 0.663 | 0.707 | 0.685 | 0.696 | 0.319 |
| PredictSNP | 100.00 | 0.678 | 0.690 | 0.704 | 0.776 | 0.738 | 0.363 |
| 3Cnet | 100.00 | 0.631 | 0.606 | 0.667 | 0.600 | 0.632 | 0.212 |

#### Data split and preprocessing

Prior to model training, the *Benchmark dataset* containing compiled predictions from the selected tools was split into training and test sets using a stratified splitting strategy to preserve the distributions of both training data independence level (*clean sequence, clean site*, and *clean variant*) and data source type. For that purpose, a composite stratification key was constructed by concatenating each variant’s training data independence level with its source type label (variants originating from ClinVar or Humsavar were designated as *Human*, while those from MMP and PMD were labeled *Mixed*). Subsequently, variants containing missing values in any of the tool prediction fields (failed predictions) were excluded and stored separately. The remaining data were split into training (80%) and test (20%) sets using the aforementioned composite stratification key, ensuring proportional representation of all independence level–source type combinations across both partitions. As raw numerical outputs from the individual VEPs span heterogeneous value ranges and directionalities, two additional steps were applied on both partitions to ensure directional alignment and comparable score ranges across all tools: (1) sign-flipping of scores negatively correlated with the label, and (2) min-max rescaling with percentile clipping to transform scores into the unified range. Specifically, per-feature lower and upper bounds were defined as the 1st and 99th percentiles of the post-flip training scores, preventing extreme outliers from compressing potentially informative score ranges. At inference, values falling outside the percentile range were hard-clamped to the output range, preventing out-of-distribution tool scores from producing extrapolated values. The preprocessed training set was further divided into training and validation subsets during classifier training via cross-validation (see Model training). The training and test sets are available at https://doi.org/10.5281/zenodo.23597130.

#### Model training

Several families of supervised machine learning-based classifiers were evaluated for their capability to generate consensus variant effect predictions from the scores of individual VEPs. These comprised logistic regression (LogReg; *sklearn.linear_model.LogisticRegressionCV*), a random forest (RF; *sklearn.ensemble.RandomForestClassifier*), an XGBoost classifier (XGB; *xgboost.XGBClassifier*), a support vector machine (SVM; *sklearn.svm.SVC*), and a multi-layer perceptron (MLP; *sklearn.neural_network.MLPClassifier*). All ten machine learning-based classifiers were trained and evaluated on the fully preprocessed training and validation sets, respectively, using 5-fold cross-validation (CV) with AUC as the target metric for maximization. Exhaustive hyperparameter optimization was performed via grid search (*sklearn.model_selection.GridSearchCV*) for the random forest, XGBoost, SVM, and MLP classifiers, and via the internal cross-validated regularization path search of LogisticRegressionCV for the logistic regression models (the complete hyperparameter grids evaluated for each classifier are provided in **Table S5**). For GridSearchCV-wrapped models, the mean test-fold AUC of the best hyperparameter configuration was used as the representative CV AUC, alongside the mean train-fold AUC from the same configuration, enabling direct assessment of the train–CV gap as a proxy for overfitting. For LogisticRegressionCV, the CV AUC was extracted directly from the internal per-fold score array, while the training AUC was computed post hoc from predictions on the full training set. All models were ranked by mean CV AUC, and the highest-ranked model was selected as the final consensus classifier. Additionally, two untrained baseline ensembles were considered in the evaluation: simple averaging of individual predictor scores, and majority voting based on the proportion of individual tools predicting a pathogenic effect, using per-tool classification thresholds derived by maximizing Youden’s index on the training set partition.

#### Model validation

Individual tools, baseline ensembles, and the selected machine learning-based classifier (XGBoost) were subsequently evaluated across both training and the held-out test set partitions with AUC as the primary metric. Uncertainty was quantified via stratified bootstrapping (N = 1000 iterations), performed separately for each evaluation partition. Each resample was drawn with replacement while preserving the positive-to-negative class ratio. The bootstrap standard deviation across 1000 estimates was reported as the uncertainty measure for each metric.

#### Feature importance analysis

The contribution of the individual tools to the consensus predictions of the model was assessed using two complementary methods: TreeSHAP and Permutation feature importance (PFI). TreeSHAP was computed using *shap.TreeExplainer*, which exploits the additive tree structure of XGBoost to compute exact Shapley values operating on the model’s raw margin (log-odds) output, prior to sigmoid transformation. This ensures mathematical exactness of the additive feature attribution, which would be lost if SHAP values were computed directly in probability space. The primary importance metric was the mean absolute SHAP value per feature, representing the average magnitude of each tool’s contribution to the model’s log-odds output across all variants in the dataset, directly comparable across tools due to the prior preprocessing of all features. Uncertainty in mean absolute SHAP values was quantified via bootstrap resampling (n=200 iterations): at each iteration, a sample of equal size was drawn with replacement, SHAP values were recomputed using TreeExplainer, and per-feature mean absolute values were recorded; the standard deviation across bootstrap iterations was reported as the uncertainty estimate. PFI was computed using a custom implementation with AUC as the scoring metric and 30 permutation repeats per feature. For each feature and each repeat, the column was randomly shuffled while all other features were held fixed; the drop in AUC relative to the unshuffled baseline was recorded as the importance score for that repeat. The mean AUC drop and its standard deviation across the 30 repeats were reported. Both analyses were conducted independently on both preprocessed training and test sets, enabling assessment of consistency between in-sample and out-of-sample importance rankings.

## Results and Discussion

### Overview of included variant effect predictors

Our literature search identified 18 candidate protein sequence-based variant effect predictors (VEPs), which were evaluated against criteria enabling robust, controlled benchmarking. In total, 11 VEPs met all criteria, such as local installation option, its training data availability (for supervised methods), and user-defined sequences as input, either by default or after minor modifications of the source code (**Table S1**). The remaining seven tools were excluded, most commonly due to unavailable model weights or training data (AlphaMissense^20^, PrimateAI^51^), unavailable or incomplete source code for the full computational pipeline (VARITY^52^, Envision^53^, DEOGEN2^54^) at the time of assessment, or extensive computational demands (mutationTCN^55^, MTBAN^56^), preventing their streamlined application to the *Benchmark dataset* (**Table S1**). Three widely used legacy tools (PROVEAN^24^, PolyPhen-2^27^, PredictSNP^36^) were additionally included as baseline comparators, reflecting their established role as community reference standards, yielding a final set of 14 VEPs installed and configured for benchmarking (**Table 1, S3, S4**).

The final set of VEPs spans a broad range of computational strategies, from feature-driven machine learning-based classifiers (e.g., PolyPhen-2, 3Cnet) to unsupervised evolutionary models built on multiple sequence alignments (e.g., GEMME, PHACT), modern protein language model embeddings (e.g., ESM1b, VESPA), and consensus-based (PredictSNP) approaches. This methodological diversity, spanning two decades of development, allowed us to evaluate not only absolute predictive performance but also whether architectural and data-source differences translate into systematic performance gains.

### Construction of benchmark dataset

Reliable benchmarking requires a comprehensive and representative leakage-controlled validation dataset. To this end, we constructed the *Benchmark dataset* by integrating data from four independent sources, systematically removing overlaps with the training data of the tested supervised predictors, and rebalancing the resulting entries across substitution types, pathogenicity classes, and data sources (**Figure 1**).

**Figure 1:**
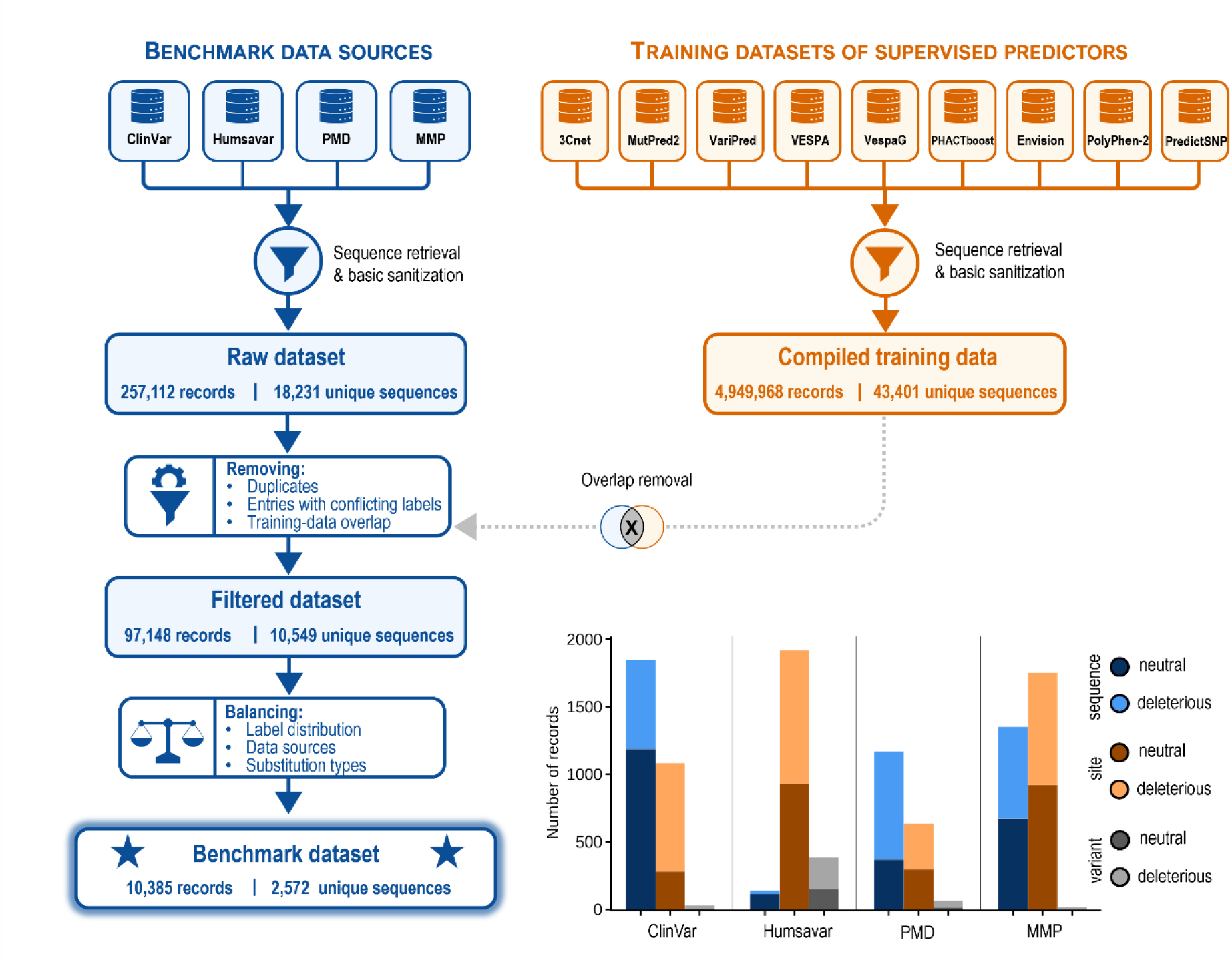
Construction and composition of the Benchmark dataset. Variant records from four independent sources: ClinVar, Humsavar, PMD, and MMP were aggregated into the *Raw dataset* following sequence retrieval and basic sanitization. Training datasets of included supervised variant effect predictors were compiled in parallel using the same retrieval procedure. The *Raw dataset* was then filtered by removing duplicate entries, records with conflicting labels, and any record overlapping with the compiled training data, forming the *Filtered dataset*. Additionally, the records in *Filtered dataset* were annotated with data independence level: i) *clean sequence* (i.e., the variant’s protein sequence was entirely absent from the training data); (ii) *clean site* (i.e., the protein sequence was present in the training data but the mutation site was not), and (iii) *clean variant* (i.e., only the specific substitution at an otherwise-seen site was absent in the training data). This dataset was subsequently rebalanced across class labels, data sources, and substitution types, producing the final *Benchmark dataset*. The bar chart describes the composition of the final *Benchmark dataset* by data source, training-data independence level, and pathogenicity label. For each source (ClinVar, Humsavar, PMD, MMP), record counts are shown stratified by independence level (*clean sequence, clean site*, and *clean variant*) and further subdivided by binary label (neutral vs. deleterious), indicated by darker and lighter shading, respectively, within each independence-level color.

Our approach differs substantially from conventionally used datasets, which typically rely solely on clinical data^35^. While these sources, such as ClinVar^40,41^ or HGMD^43^, undeniably provide the most clinically relevant data, it has been previously systematically observed that several supervised VEPs performed substantially better on clinical-observation benchmarks than on DMS-derived functional data^37,38,57^, pointing to concerns about residual circularity and potential overestimation of the performance. Therefore, we compiled our benchmark from four independent sources: ClinVar and Humsavar served as sources of clinically relevant human variants, while two additional datasets, the Protein Mutant Database^46^ (PMD) and Massively Mutated Proteins^36,47–49^ (MMP), previously compiled and used for independent validation of the original PredictSNP, provide labels derived from experimental data assessing the mutational impact on protein function, stability, or disease association across proteins from various species, collectively providing broader, cross-species representation of variant effects beyond human clinical annotations, thereby serving as a more challenging prediction target for the tested VEPs.

Combining these sources yielded the *Raw dataset* of 257,112 records spanning 18,231 unique protein sequences. After removing duplicates, conflicting labels, and variants overlapping with the training data of the supervised predictors, 97,148 records across 10,549 unique sequences remained in the *Filtered dataset*. To further probe the residual circularity, the remaining records were categorized into three data independence levels (*clean sequence, clean site*, and *clean variant*; see Materials and methods: Benchmark dataset construction), enabling direct assessment of performance degradation of the benchmarked tools as a function of independence stringency. At this stage, the dataset, however, displayed a pronounced imbalance across several attributes (**Table S2**): neutral variants outnumbered deleterious ones (67,905 vs. 29,243), ClinVar dominated the data sources (77,061 of 97,148 records), and some substitutions were heavily overrepresented (e.g., Arg→Gln, Ala→Thr) while others were rare (e.g., Cys→Thr, Cys→Ile; **Figure S1**). To correct these imbalances, we rebalanced the *Filtered dataset*, yielding the final *Benchmark dataset* of 10,385 variants from 2,572 unique sequences with a near-even distribution of labels (47% neutral, 53% deleterious) and sources (ClinVar 29%, Humsavar 24%, PMD 18%, MMP 30%), and with substantially reduced substitution-type bias (**Figure S1**). Most variants remained strongly independent of training data (43% *clean sequence*, 52% *clean site*, 5% *clean variant*; detailed composition of the dataset at each stage is provided in **Table S2**), further supporting the dataset’s suitability for unbiased VEP evaluation.

### Benchmarking of variant effect predictors

We evaluated the predictive performance of the 14 selected VEPs. Predictive performance was assessed using in-house-collected prediction scores for the constructed *Benchmark dataset*, ensuring full control over the evaluation process rather than relying on precomputed scores deposited in external databases (**Table 2**).

Most tested tools produced predictions for nearly all benchmark variants, with the exception of substitutions to selenocysteine, which could not be processed by most tools except 3Cnet, PROVEAN, and PredictSNP. The highest failure rates were observed for EVE, PolyPhen-2, and VESPA. For EVE and VESPA, failures stemmed predominantly from high computational demands on longer protein sequences, whereas PolyPhen-2 failed on proteins with an insufficient population of homologous sequences, resulting in low-quality multiple sequence alignments (MSAs) and consequently failed predictions. A similar, though less frequent, pattern was observed for GEMME, PHACT, and PHACTboost, all of which rely on MSAs. The sensitivity of these three tools to alignment quality was examined in more detail (see **Supplementary Note 1** and **Table S6**): larger, more diverse MSAs generally improved prediction accuracy for GEMME and PHACT, but at the cost of substantially increased computational time and a higher rate of failed predictions.

Overall performance across the tested predictors, assessed by AUC on the common subset (**Table 2**), was largely comparable, with values ranging from 0.631 to 0.758. Notably, ranking among the top four for VESPA and GEMME was consistent with their systematically high ranking in previously published benchmarks^57,58^. Furthermore, the best performing tools spanned distinct methodological categories: VariPred and VESPA combine protein language model (pLM) embeddings with supervised learning, whereas GEMME and PHACT are purely evolutionary-based, indicating that no single methodological paradigm consistently dominated performance on this independent benchmark. Remarkably, PROVEAN, PolyPhen-2, and PredictSNP, despite being considerably older and relying on comparatively simpler methodology and training procedures relative to more recent tools, achieved AUC values broadly comparable to several newer, more sophisticated predictors, suggesting that methodological complexity alone does not guarantee superior generalization on an independent, circularity-controlled benchmark. Threshold-dependent metrics largely followed the AUC ranking, with PHACT, VariPred, and VESPA also achieving the highest MCC values (0.395, 0.393, and 0.382, respectively). Several tools showed a marked precision-recall imbalance: PROVEAN, PHACTboost, and PredictSNP exhibited comparatively high recall (0.802, 0.749, and 0.776) but lower precision (0.686, 0.702, and 0.704), suggesting a tendency toward over-prediction of deleterious variants, whereas VariPred, PHACT, and ESM1b showed the opposite trend (precision: 0.767, 0.761, and 0.761; recall: 0.649, 0.669, and 0.630, respectively). Evaluating each tool on all of its own calculated records (**Table S7**) gave similar rankings and systematically slightly higher AUC (by 0.000– 0.027) and MCC, suggesting that the records some tools failed to predict were generally easier for the remaining tools.

Stratifying performance by data source revealed a consistent pattern across nearly all tools: AUC values were markedly higher for the two clinically annotated human datasets (ClinVar, Humsavar) than for the two functional, cross-species datasets (PMD, MMP) (**Table S8**). This gap was most pronounced for 3Cnet, whose AUC dropped from 0.798 on ClinVar to 0.498 on PMD and 0.526 on MMP, respectively, close to the values for random classification, suggesting a strong dependence on features or training data specific to human clinical variants that fails to generalize to functional, non-human variant-effect data. The dramatic drop in 3Cnet’s performance on PMD and MMP data partially explains its overall lower ranking in our full benchmark despite ranking among the top-performing predictors in other published benchmarks^30,35^, which were conducted on clinically annotated human data. A similar, although less extreme, pattern was observed for VariPred, MutPred2, and ESM1b, each showing a 0.20-0.25 AUC decrease from ClinVar to PMD/MMP. In contrast, VespaG and VESPAl showed comparatively balanced performance across sources (e.g., VespaG: 0.774 on ClinVar versus 0.734 on both PMD and MMP), indicating better generalization beyond human clinical variants. These findings align with previous reports of higher tool performance on clinical benchmarks relative to deep mutational scanning datasets^58^. This effect was pronounced the most in supervised tools, in line with previous studies^37^, although we observed similar trend for most of the tools, including unsupervised ESM1b.

Stratification by training-data independence level revealed a similar but weaker pattern, with AUC generally highest in the least strict *clean variant* stratum compared with the *clean site* and *clean sequence* strata (**Table S8**). However, this difference was markedly smaller than the drop observed across data sources, and the comparatively small size of the *clean variant* stratum (N = 434), together with its skew toward Humsavar-derived variants (Bar plot in **Figure 1**), precludes strong conclusions about residual circularity from this analysis alone. Nevertheless, our results indicate that the performance of most tested VEPs, especially supervised methods such as 3Cnet and VariPred, depends strongly on the data source and partially on the independence level from their training data, which may lead to inflated apparent performance. Notably, while type 1 circularity, represented by direct overlap of training and testing datapoints, was thoroughly addressed in our benchmark by removal of the overlapping records, residual leakage, particularly type 2 circularity, may persist despite being partially controlled through the independence annotations. Type 2 circularity, characterized by shared features between training and testing records (such as variants of the same or similar proteins), is harder to control, since even unsupervised and pLM-based models may have been trained on sequences that cannot be explicitly identified or excluded, and our stratification did not account for near-identical homologs. To probe this limitation, we tested tool performance on a subset of the *clean sequence* stratum filtered to exclude close homologs of the compiled training data (methodology and results are detailed in **Supplementary Note 2**). The performance of most of the tools was essentially unchanged or only marginally lower with differences within one to two bootstrap standard deviations of the AUC estimate (SD ≈ 0.02) on this subset. Larger decreases (0.06–0.07, about three standard deviations) occurred only for ESM1b, 3Cnet, and VespaG (**Table S9**). In the case of 3Cnet and VespaG, these results may suggest that part of their performance reflects homology to training or pretraining sequences. However, given the small size of the dataset (N = 658), these results are indicative only, and larger datasets and focused analyses are needed to confirm which tools are significantly affected by homology-based circularity. Overall, our results indicate that homology-based circularity may affect some tools but does not appear to be the dominant driver of the inflated performance observed overall.

### Consensus model development and validation

Motivated by the relative underperformance of our earlier PredictSNP meta-predictor compared to the majority of recently developed VEPs, we developed a novel consensus model, PredictSNP^prot^, which combines predictions from a set of validated tools into a single, robust variant-effect prediction score, serving as an improved successor to the original PredictSNP. Only tools exhibiting low failure rate (< 2%) on the *Benchmark dataset*, and those capable of efficient scaling to complete saturation mutagenesis were retained as constituents of the model (3Cnet, ESM1b, VESPAl, VespaG, PROVEAN, VariPred, PHACT, PHACTboost, and GEMME). The remaining five tools were excluded from the pool: PredictSNP (superseded by this consensus model), PolyPhen-2, EVE, and VESPA (observed failure rate > 2%), and MutPred2 (unable to scale to predict full saturation mutagenesis).

A diverse set of machine learning models, comprising L1-regularized logistic regression (LogReg_L1), elastic net logistic regression (LogReg_EN), two hyperparameter configurations of Random Forest (RF_1, RF_2), XGBoost (XGB_1, XGB_2), Support Vector Machine (SVM_1, SVM_2), and Multilayer Perceptron (MLP_1, MLP_2), alongside two baselines (majority voting and mean score) was evaluated to identify the optimal strategy for aggregating individual tool scores into a single pathogenicity prediction. All models were trained and evaluated via cross-validation using exhaustive hyperparameter optimization (**Table S5**), yielding the best-performing configuration per model family (**Table 3**).

**Table 3:** Cross-validation performance of evaluated models. AUC = area under the receiver operating characteristic curve. SD = standard deviation. CV = 5-fold cross-validation; CV — Train and CV — Val. report mean AUC and SD across folds on the training and held-out validation folds, respectively. NA = not applicable. L1 = L1 (Lasso) regularization; EN = elastic net regularization. The best model is highlighted in bold and denoted by ✓. Indexes in model names indicate the hyperparameter grid configuration for a specific model.

| **Category** | **Model** | **CV — Train** |  | **CV — Val.** |  |
|---|---|---|---|---|---|
|  |  | **AUC** | **SD** | **AUC** | **SD** |
| **ML-based** | **XGB_2 ✓** | **0.890** | **0.002** | **0.826** | **0.009** |
|  | XGB_1 | 0.866 | 0.002 | 0.825 | 0.009 |
|  | RF_2 | 0.860 | 0.002 | 0.822 | 0.009 |
|  | RF_1 | 0.864 | 0.002 | 0.821 | 0.009 |
|  | MLP_2 | 0.825 | 0.003 | 0.817 | 0.011 |
|  | MLP_1 | 0.818 | 0.002 | 0.813 | 0.009 |
|  | SVM_1 | 0.814 | 0.003 | 0.807 | 0.009 |
|  | SVM_2 | 0.841 | 0.003 | 0.806 | 0.010 |
|  | LogReg_EN | 0.800 | NA | 0.799 | 0.007 |
|  | LogReg_L1 | 0.800 | NA | 0.799 | 0.007 |
| **Baseline ensembles** | Mean score | 0.793 | 0.003 | 0.793 | 0.012 |
|  | Majority voting | 0.794 | 0.003 | 0.790 | 0.013 |

Machine learning models consistently outperformed both baseline ensembles, demonstrating the added value of a learned consensus over simple heuristic approaches. All tested models exhibited comparable validation AUC with XGBoost and Random Forest achieving the highest validation AUC, of 0.826 ± 0.009 and 0.822 ± 0.009, respectively. Although several of XGBoost’s optimal hyperparameter values lay at the boundaries of the initial search grid (XGB_2 in **Table S10**), an extended grid search (XGB_3 in **Table S10**) yielded only marginal gains at the cost of increased overfitting (**Table S11**), indicating that the initial configuration had already reached the effective performance ceiling for this training dataset. XGB_2 was therefore selected as the final PredictSNP^prot^ model based on highest validation AUC from the initially tested models. Its generalization performance was subsequently benchmarked against the individual tools and baseline ensembles on a held-out test set comprising 20% of the full *Benchmark dataset*, stratified to preserve the distributions of training-data independence level and data source (**Table 4**).

**Table 4:** Performance of PredictSNP^prot^, baseline ensembles and individual VEPs on the held-out test set. AUC = area under the receiver operating characteristic curve. SD = standard deviation. Standard deviations were estimated from bootstrapping (N = 1000). PredictSNP^prot^ corresponds to the XGB_2 model, identified previously using cross-validation. Training partition (N = 8,144) refers to the partition of the Benchmark dataset used to train the consensus model; individual VEPs were not trained on this partition. Test set (N = 2,037) refers to the held-out partition of the Benchmark dataset used for independent validation of the model.

| Category | Model | Trainset |  | Test set |  |
|---|---|---|---|---|---|
|  |  | AUC | SD | AUC | SD |
| **ML-based** | **PredictSNP<sup>prot</sup>** | **0.883** | **0.004** | **0.818** | **0.010** |
| **Baseline ensembles** | Mean score | 0.793 | 0.005 | 0.772 | 0.011 |
|  | Majority voting | 0.794 | 0.005 | 0.770 | 0.011 |
| **Individual tools** | VariPred | 0.772 | 0.005 | 0.761 | 0.011 |
|  | VESPA* | 0.763 | 0.005 | 0.752 | 0.011 |
|  | PHACT | 0.754 | 0.005 | 0.737 | 0.011 |
|  | PolyPhen-2* | 0.726 | 0.006 | 0.736 | 0.012 |
|  | VESPAl | 0.738 | 0.006 | 0.735 | 0.011 |
|  | GEMME | 0.756 | 0.005 | 0.733 | 0.012 |
|  | MutPred2 | 0.749 | 0.006 | 0.730 | 0.012 |
|  | PROVEAN | 0.725 | 0.006 | 0.722 | 0.012 |
|  | VespaG | 0.739 | 0.005 | 0.720 | 0.012 |
|  | ESM1b | 0.744 | 0.005 | 0.715 | 0.012 |
|  | PHACTboost | 0.728 | 0.006 | 0.708 | 0.012 |
|  | EVE* | 0.702 | 0.006 | 0.696 | 0.013 |
|  | PredictSNP | 0.684 | 0.005 | 0.682 | 0.010 |
|  | 3Cnet | 0.658 | 0.006 | 0.640 | 0.012 |

An asterisk (\*) indicates that these tools were evaluated on the datapoints, for which they produced a prediction (VESPA: N<sub>train</sub> = 8,049, N<sub>test</sub> = 2,005; PolyPhen-2: N<sub>train</sub> = 7,869, N<sub>test</sub> = 1,973; EVE: N<sub>train</sub> = 7,121, N<sub>test</sub> = 1,743; other tools, ensembles and PredictSNP<sup>prot</sup> scored all records from the respective datasets.)

On the held-out test set, PredictSNP^prot^ achieved an AUC of 0.818 ± 0.010, a modest ∼0.01 decrease relative to its cross-validation estimate, demonstrating strong generalization and robustness. Moreover, it outperformed all individual evaluated VEPs, exceeding the best-performing constituent tool, VariPred (AUC = 0.761), by approximately 5.7 percentage points, further confirming that the learned consensus meaningfully improves predictive performance over the individual tools. Notably, our model showed striking improvements compared to the original PredictSNP, surpassing its test-set AUC by 13.6 percentage points and demonstrating that the meta-prediction discriminative power heavily depends on the quality and diversity of its constituent tools.

Having established the superior discriminative performance and generalization of PredictSNP^prot^, we next examined the relative contribution of each constituent tool to the consensus prediction (**Figure 2, Table S12**). Feature contribution analysis revealed a consistent hierarchy of tool importance within the PredictSNP^prot^ consensus model, with VariPred emerging as the dominant contributor by a substantial margin, ranking first by both treeSHAP and PFI on both the training and test sets (mean |SHAP| = 0.581 ± 0.004 train, 0.584 ± 0.008 test; PFI ΔAUC = 0.067 ± 0.003 train, 0.057 ± 0.006 test), consistent with its status as the best-performing individual tool in the benchmarking (**Table 2**). PHACT followed as the second-ranked contributor by both methods on both splits, also in line with its comparatively strong standalone performance. Notably, 3Cnet ranked third in feature importance despite being the weakest-performing individual tool in the benchmark (test AUC = 0.640, the lowest among all 14 evaluated VEPs), indicating that its predictive signal is beneficially orthogonal to that of the higher-performing tools and is thus enhanced by the consensus model relative to its standalone accuracy. XGBoost, therefore, identified and exploited this complementary, non-redundant information rather than simply weighting tools in proportion to their standalone performance. This capacity to extract orthogonal signals from otherwise low-ranking tools likely contributes to superior test-set performance of PredictSNP^prot^ relative to its best single constituent (**Table 4**). Finally, the close agreement between treeSHAP and PFI rankings, and between training and test partitions, indicates that the model’s reliance on individual tool scores is stable and not an artifact of overfitting.

**Figure 2:**
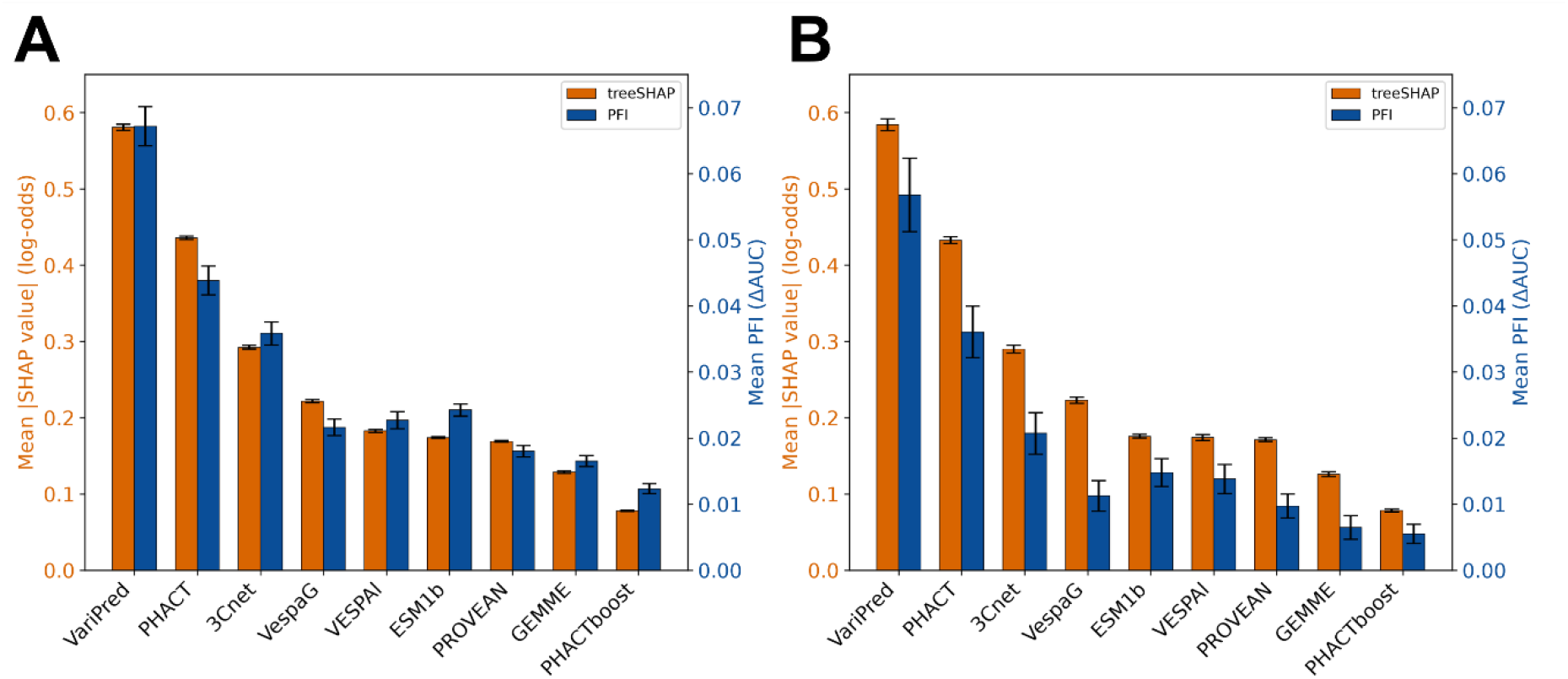
Contribution of individual VEP scores to PredictSNP^prot^consensus predictions on the training (A) and test (B) sets. Orange bars (left axis) show the mean absolute treeSHAP value per feature, with error bars denoting the bootstrap standard deviation across 200 resamples. Blue bars (right axis) show the mean permutation feature importance (PFI; the average drop in AUC when a feature’s values are independently shuffled across 30 repeats, relative to the unpermuted baseline) with error bars denoting the standard deviation across these repeats.

## Conclusions

In this study, we performed a comprehensive, independent benchmark of state-of-the-art variant effect predictors (VEPs). We assembled a new *Benchmark dataset* combining human and cross-species variants from four independent sources and refined it to remove any overlap with the training data of all evaluated supervised tools, enabling fair, leakage-controlled evaluation. Our initial review of available predictors showed that only a fraction of published VEPs can be independently reproduced or fairly benchmarked. In many cases, model weights, complete source code, or training data were inaccessible, preventing independent installation, prediction, and performance assessment. Using these criteria, we identified 11 VEPs spanning a wide range of underlying strategies, from classical alignment-based scoring to protein language models and phylogeny-aware architectures. Furthermore, the initial pool of selected VEPs, representing recent state-of-the-art methods, was extended by 2 older, widely adopted tools and our previously developed meta-predictor, PredictSNP. The benchmark revealed a relatively narrow performance range (AUC 0.631–0.758) across all tested tools, with VariPred and VESPA ranking the highest. Notably, even older, methodologically simpler tools such as

PROVEAN and PolyPhen-2, achieved performance broadly comparable to newer, more computationally sophisticated predictors, underscoring the challenging nature of further performance improvements in the field.

Building on this benchmark, we developed PredictSNP^prot^, a gradient-boosting-based consensus model integrating nine of the validated tools. The model achieved a held-out test-set AUC of 0.818 ± 0.010, outperforming its best individual constituent, VariPred, by 5.7 percentage points and the older PredictSNP by striking 13.6 percentage points. Feature contribution analysis confirmed that this improvement does not simply reflect favoring the strongest individual tool, but rather leveraging tool complementarity in making predictions. These results establish PredictSNP^prot^ as a robust, highly accurate successor to the original PredictSNP. More broadly, this work provides the community with a fair, reproducible benchmark of state-of-the-art VEPs (the *Benchmark dataset* including the compiled predictions is available at https://doi.org/10.5281/zenodo.23597130. Furthermore, it introduces a new consensus model, PredictSNP^prot^ (available at https://github.com/loschmidt/PredictSNPprot), which will be further integrated into the PredictSNP webserver, delivering accurate variant effect predictions for both clinical variant interpretation and protein engineering applications.

## Acknowledgments

The authors would like to express their thanks to the European Union’s Horizon 2020 research and innovation programme under grant agreement No. 857560 (CETOCOEN Excellence), Horizon Europe Framework programme under grant agreement No. 101136607 (CLARA) and co-funded by the EU from the Operational Programme Jan Amos Komenský (no. CZ.02.01.01/00/23_029/0008437), Grant Agency of the Czech Republic (No. 25-18233M) and the Ministry of Health of the Czech Republic in cooperation with the Czech Health Research Council within the National Institute for Cancer Research (No. NW26A-NUVR). Computational resources were provided by the e-INFRA CZ, ELIXIR-CZ, and RECETOX RI projects (90254, LM2023055, and LM2023069), supported by the Ministry of Education, Youth and Sports of the Czech Republic. Views and opinions expressed are however those of the author(s) only and do not necessarily reflect those of the European Union or REA. Neither the European Union nor the granting authority can be held responsible for any use that may be made of the information it contains. MH is supported by the scholarship Ph.D. Talent funded by the Brno City Municipality.

## Author contributions

M.H. and P.K. contributed equally to this work. M.H. selected the variant effect predictors included in the study, collected and curated the datasets, and performed the benchmarking analyses. P.K. installed and configured the predictors. M.H. and P.K. developed the consensus model and conducted the analysis and interpretation of results. M.H. and P.K. drafted the manuscript, with input from all authors. M.M., S.M., D.B., and J.D. supervised the research. D.B. and J.D. conceived and designed the study. All authors reviewed and approved the final manuscript.

## Data and code availability

Supplementary data is available in Supplementary Information. The datasets generated in this study, including all stages of *Benchmark dataset* construction, compiled predictions of the benchmarked tools, and dataset splits used for training and testing of PredictSNP^prot^ are publicly available in Zenodo public repository at https://doi.org/10.5281/zenodo.23597130.

All code used for PredictSNP^prot^ development, including scripts for training PredictSNP^prot^ and for generating predictions given the scores of the constituent tools, is available on GitHub at https://github.com/loschmidt/PredictSNPprot. The repository includes the model training scripts, along with the trained model weights and parameters (JSON file) required to reproduce PredictSNP^prot^ predictions from a dataset of compiled input features. Modified versions of third-party variant effect predictors are available in separate, individually linked repositories (see **Table S3** for details).

