## Supplementary Information for "Leakage-Controlled Benchmarking of Variant Effect Predictors and Development of Gradient-Boosting Consensus Model"

### Supplementary material

#### Supplementary Notes

##### Supplementary Note 1: Details on effect of multiple sequence alignment (MSA) on performance of GEMME, PHACT, and PHACTboost

Since GEMME[1], PHACT[2], and PHACTboost[3] rely on an MSA to compute mutational effect scores, we examined the sensitivity of their predictions on the input MSA. For PHACT and PHACTboost, MSA generation is by default integrated directly into their shared pipeline, comprising a sequence search via PSI-BLAST[4], hit filtering, and alignment using MAFFT[5] (FFT-NS-1), whereas GEMME requires a pre-built MSA as input. To enable efficient, consistent predictions across all three tools, the MSA-generation step from the PHACT pipeline was externalized and used as a single, unified preprocessing stage to generate MSA input for all three predictors (see the MSA pipeline in **Table S3** and **Table S4**).

This externalized workflow was further optimized to maximize predictive performance while minimizing failure rate, using the Benchmark dataset (2,572 unique protein sequences; 10,385 variant entries). Two processing steps were evaluated independently: (i) a filtering step, inherited from the original PHACT workflow, that limits the number of BLAST-identified hits to 1,000 and retains only sequences with 30–95% identity to the query and an E-value below 1e-5; and (ii) a clustering step (MMseqs2 easy-cluster[6,7]: min-seq-id 0.9, cov-mode 1, coverage 0.8, cluster-mode 2) that reduces MSA size and redundancy while preserving sequence diversity. Combining the presence or absence of these two steps yielded four MSA-generation configurations, each producing alignments of different size and diversity, which were evaluated for their effect on the prediction performance of GEMME, PHACT, and PHACTboost (**Table S6**).

For each configuration, MSA size (number of aligned sequences) and mean identity to the query sequence were computed per protein by comparing every aligned sequence position-by-position against the query at all non-gap query columns, expressed as the percentage of matching residues over query length, and averaged across all proteins in the benchmark dataset. Filtering substantially increased mean identity to the query (from 44% to 71% for full MSAs, and from 40% to 70% for clustered MSAs) by removing distant BLAST hits, while clustering primarily reduced MSA size and redundancy (from a mean of 10,227 to 2,680 sequences without filtering, and from 970 to 179 with filtering) without markedly affecting mean identity.

The results revealed that tool performance is generally sensitive to the composition of the input MSA, most notably for GEMME, whose AUC ranged from 0.710 with MSAs derived from clustered-filtered sequences to 0.761 with MSAs from full-unfiltered sequences, indicating that larger, more diverse MSAs improve GEMME's predictions as such MSAs contain richer information for conservation-based analyses[8]. A similar but less pronounced trend was observed for PHACT (AUC 0.764 and 0.730 for full-unfiltered and clustered-filtered MSAs, respectively). However, PHACT's failure rate increased substantially with larger MSAs, reflected in a drop in the fraction of successfully scored variants from 98.5% (full-filtered) to 93.4% (full-unfiltered), driven by higher computational demands of processing larger alignments. PHACTboost achieved its best performance with MSAs from full-filtered and clustered-unfiltered sequences, while maintaining a relatively consistent success rate (98.2–98.6%) across all four configurations.

Based on these results, the clustered-unfiltered configuration was selected as the optimal workflow, as it achieved near-best performance across all three predictors while keeping computational cost and failure rate low. The complete parametrization of the final MSA-generation procedure is detailed in **Table S3** and **S4** (MSA pipeline).

**Supplementary Note 2: Homology-Based Filtering of the Clean-Sequence Benchmark**

Our independence-level stratification (clean sequence, clean site, clean variant) filters benchmark variants based on whether their exact protein sequence, site, or variant is present in each tool's compiled training data. This criterion addresses direct data leakage (type 1 circularity) but does not account for near-identical homologous sequences that were not explicitly present in training data yet may still confer an unfair performance advantage. To evaluate whether such residual homology inflates apparent performance, we constructed a homology-filtered subset of the clean-sequence stratum and re-evaluated all tools on this subset.

Unique protein sequences from the clean-sequence benchmark stratum and from the compiled training data were extracted, deduplicated, and searched against one another using MMseqs2 easy-search[7]. A benchmark sequence was flagged as failing the homology filter if any training sequence satisfied all of the following criteria simultaneously: sequence identity ≥ 30%, both query and target alignment coverage ≥ 80% (coverage mode 0), and E-value $\leq 1 \times 10^{-3}$ (search sensitivity 7.5, alignment mode 3). Benchmark sequences without any qualifying training-sequence hit were retained in the homology-filtered subset; all others were excluded. Flags assigned at the unique-sequence level were propagated back to all corresponding benchmark variant rows before recomputing tool performance.

Applying this filter retained approximately 18% of the clean-sequence stratum (814 of 4,502 variants), indicating that the majority of clean-sequence entries share substantial similarity with at least one training sequence despite lacking an identical match. From this subset, only 658 records were scored by all tools, further reducing the size of the dataset. Despite this substantial reduction in sample size, AUC, measured on the subset, where all tested tools produced a prediction, decreased by 0.01–0.04 for eight tools and was essentially unchanged ($|\Delta AUC| \leq 0.006$) for PHACT, GEMME, and PHACTboost. For most tools these differences were within one to two bootstrap standard deviations of the AUC estimate ($SD \approx 0.02$) and should therefore not be over-interpreted. Larger decreases (0.06–0.07, about three standard deviations) occurred only for ESM1b, 3Cnet, and VespaG (**Table S9**). While for 3Cnet and VespaG, these results suggest that part of their performance reflects homology to training or pretraining sequences, similar explanation cannot be drawn for ESM1b since its pLM pretraining sets were not part of the filter (ESM1b represents zero-shot predictor, as such, it was not trained on labeled dataset). Given the small size of the homolog-free common subset (N = 658), these results are indicative only, and larger datasets and focused analyses are needed to confirm which tools are significantly affected by homology-based circularity. Overall, our results indicate that homology-based circularity may affect some tools but does not appear to be the dominant driver of the inflated performance observed overall.

#### Supplementary Figures

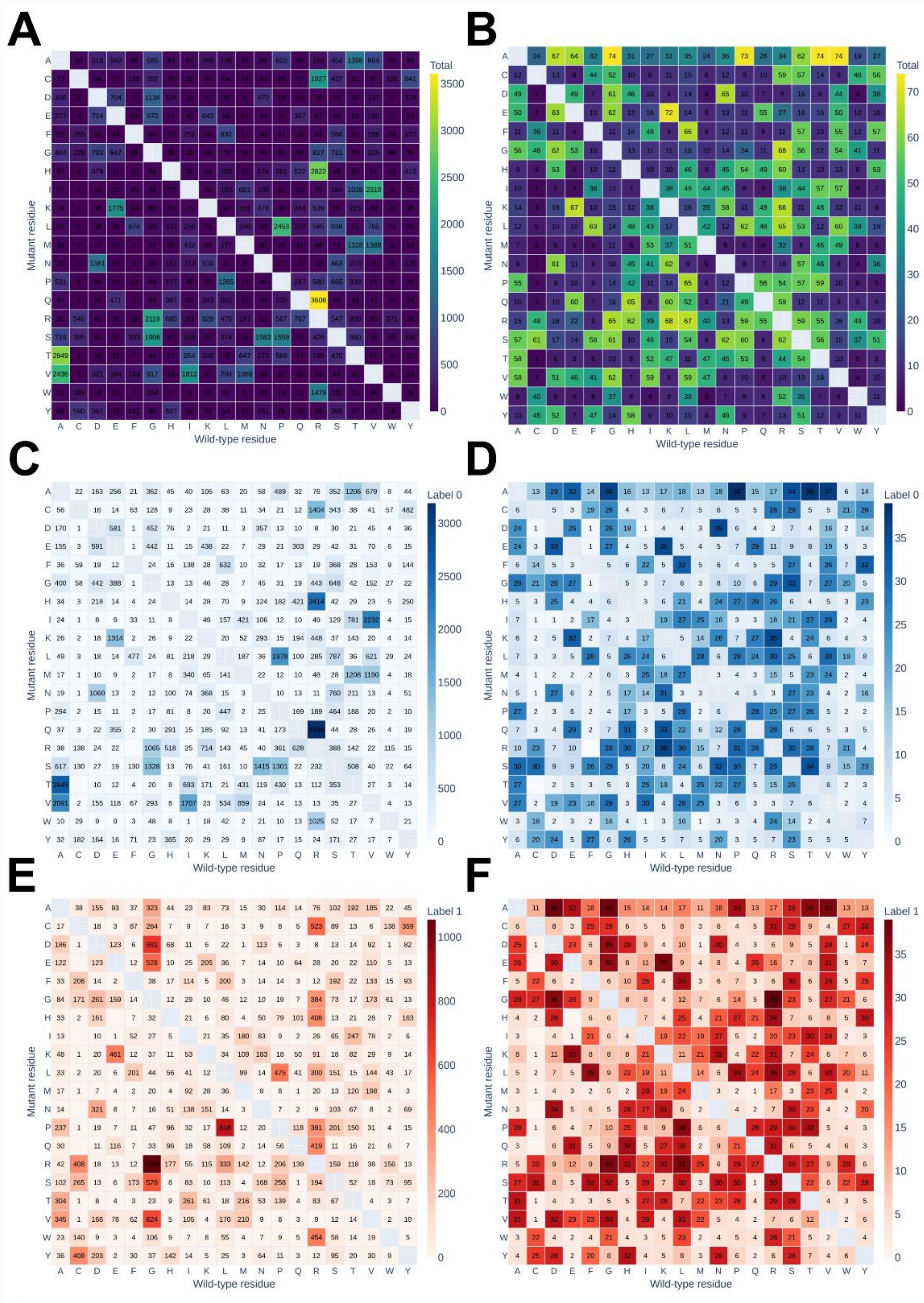


**Figure S1: Substitution-type distribution in the Filtered and Benchmark datasets.** Heatmaps show the number of records for each wild-type-to-mutant amino acid substitution pair in the *Filtered dataset* **(A, C, E)** and the final *Benchmark dataset* **(B, D, F). (A, B)** All substitutions, regardless of pathogenicity label. (C, D) Substitutions restricted to neutral variants. **(E, F)** Substitutions restricted to deleterious variants. Rows indicate the mutant residue, and columns indicate the wild-type residue; diagonal cells (no amino acid change) are masked. Color scale indicates the total record count per substitution pair, with the scale range fitted independently to each panel.

#### Supplementary Tables

**Table S1: Selection criteria applied to candidate variant effect predictors for inclusion in the benchmark.** Tools were evaluated against three sequential selection criteria: (i) availability of an installable, locally runnable version, (ii) availability of the training data (or unsupervised status, where no training data is applicable), and (iii) support for user-defined protein sequence input. "YES/NO" indicates the criterion satisfaction status based on each tool's publicly available code, model weights, and training data as of 2024; subsequent updates or releases by the original authors may change a tool's status with respect to these criteria.; "Unsupervised" indicates the tool does not require training data by design and is therefore treated as satisfying this criterion

| Tool | Installable | Training data available | User-defined sequence input | Included in final benchmark | Additional notes | Reference |
|---|---|---|---|---|---|---|
| **3Cnet** | YES | YES | YES | YES | N/A | [9] |
| **MutPred2** | YES | YES* | YES | YES | *HGMD derived training data not available | [10] |
| **ESM1b** | YES | Unsupervised | YES | YES | N/A | [11] |
| **PHACT** | YES | Unsupervised | YES* | YES | *User-defined sequence input possible upon custom changes of the source code | [2] |
| **PHACTboost** | YES | YES | YES* | YES | *User-defined sequence input possible upon custom changes of the source code | [3] |
| **VESPA** | YES | YES | YES | YES | N/A | [12] |
| **VESPAl** | YES | YES | YES | YES | N/A | [12] |
| **VespaG** | YES | YES | YES | YES | N/A | [13] |
| **GEMME** | YES | Unsupervised | YES* | YES | *Requires MSA on the input | [1] |
| **VariPred** | YES | YES | YES | YES | N/A | [14] |
| **EVE** | YES | Unsupervised | YES* | YES | *Requires MSA on the input | [15] |
| **PROVEAN** | YES | Unsupervised | YES | YES | N/A | [16] |
| **PolyPhen-2** | YES | YES | YES | YES | N/A | [17] |
| **PredictSNP** | NO* | YES | YES | YES | * accessed via webserver API | [18] |
| **MTBAN** | YES | Unsupervised | YES | NO* | *Extensive computational demands | [19] |
| **mutationTCN** | YES | Unsupervised | YES | NO* | *Extensive computational demands | [20] |
| **Alphamissense** | NO* | NO | n.t. | NO | *Model weights are not available | [21] |
| **PrimateAI** | YES | NO* | n.t. | NO | *Training data should be available through Illumina | [22] |
| **VARITY** | NO* | n.t. | n.t. | NO | *Input features calculation not included | [23] |
| **Envision** | NO* | YES | n.t. | NO | *Incomplete source code available | [24] |
| **DEOGEN2** | NO | NO | YES | NO | N/A | [25] |

An asterisk (*) denotes a qualified YES/NO for that tool, with the specific condition or caveat detailed in the Additional notes. N/A indicates that no additional clarification is needed for that entry. n.t. ("not tested") indicates the criterion was not evaluated because the tool had already been excluded at an earlier selection level. Selection proceeded sequentially: only tools satisfying Installable and Training data availability (or being unsupervised) were assessed for User-defined sequence input, and only tools satisfying all three criteria were included in the final benchmark. Tools are grouped into included recently published tools, included legacy/established tools retained for comparison, and tools excluded from the final benchmark.

**Table S2: Composition of the Benchmark dataset at each stage of construction.** The initial dataset (**Raw**) was filtered for duplicates, conflicting labels, and overlaps with training data (**Filtered**) and then rebalanced across pathogenicity labels, data sources, and substitution types (**Benchmark**, see Materials and methods). Independence level reflects each record's relationship to the training data of the evaluated supervised VEPs: clean sequence indicates the protein sequence was entirely absent from the training data; clean site indicates the protein sequence was present but the mutation site was not; clean variant indicates both the protein and mutation site were present, but not the specific substitution.

| Dataset | Total entries | Total unique seq | Label | | Source | | | | Independence level | | |
|---|---|---|---|---|---|---|---|---|---|---|---|
| | | | 0 | 1 | ClinVar | Humsavar | PMD | MMP | Sequence | Site | Variant |
| **Raw** | 257112 | 18231 | 158687 | 98425 | 169609 | 72291 | 3495 | 11717 | | N/A* | |
| **Filtered** | 97148 | 10549 | 67905 | 29243 | 77061 | 5193 | 3211 | 11683 | 12452 | 65192 | 19504 |
| **Benchmark** | 10385 | 2572 | 4931 | 5454 | 2960 | 2442 | 1864 | 3119 | 4502 | 5385 | 498 |

*N/A indicates that independence-level classification had not yet been performed at that stage of dataset construction, as this categorization is only assigned following the filtering step described in the Methods.

**Table S3: Installation and configuration of the benchmarked variant effect predictors.** Version or commit hash, model or embedding checkpoint, parameters, and local code modifications for each tool. Tools are listed in the order of Table 2. N/A, not applicable or not specified; Default, default parameters of the tool.

| Tool | Version / commit | Model / embeddings | Parameters | Code modifications / notes | Source |
|---|---|---|---|---|---|
| **VariPred** | 8b1c225 | esm2_t33_650M_UR50D | Default | Modified to reuse cached wild-type embeddings instead of recomputing them per mutant, and to output a continuous score. | github.com/loschmidt/VariPred |
| **VESPA** | 34fd1ab | Rostlab/prot_t5_xl_uniref50 | Default | None. | github.com/Rostlab/VESPA |
| **PHACT** | a8e49c8 | N/A | Default | Modified to run on any query sequence without a UniProt ID; MSA generation externalized (see MSA pipeline) and MSA processing handled upstream with author-provided scripts. Output based on the wl_param_CountNodes_3 parametrization, selected by benchmarking. | github.com/loschmidt/PHACT |
| **GEMME** | N/A | N/A | Default | MSA precomputed with the custom MSA pipeline (see below) and processed with GEMME's included script. | Docker: elodielaine/gemme@sha256:a35099f7dee9411842e3e04318f4a9b435541bce16aff1c9ef5baa507baf5e7d |
| **VESPAl** | 34fd1ab | Rostlab/prot_t5_xl_uniref50 | Default | Shares the VESPA pipeline and model checkpoint; differs only in output computation. | github.com/Rostlab/VESPA |
| **ESM1b** | 0c33758 | esm1b_t33_650M_UR50S | Default | None. | github.com/ntranoslab/esm-variants |
| **VespaG** | 3d47582 | facebook/esm2_t36_3B_UR50D | Default | None. | github.com/JSchlensok/VespaG |
| **MutPred2** | N/A | N/A | -p 1 (protein homology profiles); -c 1 (predicted conservation scores if no precomputed alignment); -b 0 (PSI-BLAST disabled if no precomputed PSSM exists); -t 0.1 (report mechanisms at P < 0.1) | None. | mutpred.mutdb.org (download) |
| **PolyPhen-2** | v2.2.3 | HumDiv-trained model | Default | Database, source code, and parameters inherited from the original PredictSNP setup. | N/A |
| **PHACTboost** | 8c100d8 | N/A | Default | Minor refactoring to prevent ID collisions. | github.com/loschmidt/PHACTboost |
| **PROVEAN** | v1.1.3 | N/A | Default | None. | jcvi.org/research/provean |
| **EVE** | 460d70e | N/A | Default | None. | github.com/OATML-Markslab/EVE |
| **3Cnet** | ddf452e | N/A | -e 36 | Modified to accept custom sequences via an added input parameter. | github.com/loschmidt/3Cnet |
| **MSA pipeline (shared by PHACT, PHACTboost, GEMME)** | N/A | N/A | PSI-BLAST: e-value 1e-5, 2 iterations, max 10,000 sequences. MMseqs2 easy-cluster: --min-seq-id 0.9, --cov-mode 1, -c 0.8, --cluster-mode 2 (greedy, CD-HIT-like). MAFFT: --anysymbol --retree 1 --maxiterate 0 (FFT-NS-1). | Shared upstream MSA-generation pipeline; database: UniRef100 (Table S4). | N/A |

**Table S4: Sequence databases used by the benchmarked tools.** Size and release date of the sequence databases used for homology search or MSA construction.

| Database | Used by | Sequences | Release date | BLAST DB format |
|---|---|---|---|---|
| nr | MutPred2 | 26,682,258 | Jun 28, 2013 | N/A |
| nr | PROVEAN | 15,006,980 | Aug 15, 2011 | v4 |
| UniRef100 (uniref100-formatted.fasta) | PolyPhen-2 | 19,211,424 | Nov 14, 2012 | v4 |
| UniRef100 | MSA pipeline (PHACT, PHACTboost, GEMME) | 475,217,233 | Apr 13, 2026 | v5 |

**Table S5: Hyperparameter grids of the consensus classifiers.** For each model, all combinations of the listed values were evaluated by exhaustive grid search with 5-fold cross-validation, using ROC-AUC as the selection metric (LogisticRegressionCV searched its regularization path internally). Parameters with a single value were held fixed. Grid size is the number of hyperparameter combinations. Cs = 20 values log-spaced between $10^{-4}$ and $10^{4}$ (scikit-learn default for an integer Cs). class_weight None denotes no class weighting. hidden_layer_sizes lists the number of units per hidden layer. Grid sizes are nominal (e.g., gamma has no effect for the linear kernel of SVM1). Parameters not listed were left at their library defaults.

| Model | Searched hyperparameters | Fixed settings | Grid size |
|---|---|---|---|
| **LogReg_L1** | Cs: 20 values, log-spaced | penalty = l1; solver = saga; l1_ratio = 1; max_iter = 10000; cv = 5 | 20 |
| **LogReg_EN** | Cs: 20 values, log-spaced; l1_ratio: 0.01, 0.05, 0.1, 0.3, 0.5, 0.7, 0.9, 0.95, 0.99 | penalty = elasticnet; solver = saga; max_iter = 10000; cv = 5 | 180 |
| **RF_1** | max_depth: 3, 5, 7; max_features: 0.3, 0.5; min_samples_leaf: 20, 50, 100; min_samples_split: 50, 100, 150; n_estimators: 50, 100, 150 | – | 162 |
| **RF_2** | ccp_alpha: 0.0, 0.001, 0.005; class_weight: balanced, balanced_subsample, None; max_depth: 3, 5, 7; max_features: 0.2, 0.3, 0.5; max_samples: 0.6, 0.7, 0.8; min_samples_leaf: 20, 50, 100; n_estimators: 100, 200, 300 | – | 2,187 |
| **XGB_1** | colsample_bytree: 0.6, 0.8; learning_rate: 0.01, 0.05; max_depth: 2, 3, 4; n_estimators: 50, 100, 150; reg_alpha: 1.0, 5.0; reg_lambda: 2.0, 5.0, 10.0; subsample: 0.6, 0.8 | – | 432 |
| **XGB_2** | colsample_bytree: 0.5, 0.6, 0.8; gamma: 0.1, 0.5, 1.0; learning_rate: 0.005, 0.01, 0.05; max_depth: 2, 3, 4; min_child_weight: 5, 10, 20; n_estimators: 100, 200, 300; reg_alpha: 1.0, 5.0, 10.0; reg_lambda: 2.0, 5.0, 10.0; subsample: 0.5, 0.6, 0.8 | – | 19,683 |
| **XGB_3** | colsample_bytree: 0.8, 0.9, 1.0; gamma: 0.05, 0.1, 0.5; learning_rate: 0.025, 0.05, 0.075, 0.1; max_depth: 3, 4, 5; min_child_weight: 3, 4, 5; n_estimators: 300, 350, 400; reg_alpha: 0.5, 1.0, 1.5; reg_lambda: 2.0, 5.0, 7.5, 10.0, 12.5; subsample: 0.6, 0.8, 0.9 | – | 43,740 |
| **SVM_1** | C: 0.1, 1, 10, 100; gamma: scale, auto, 0.001, 0.01, 0.1; kernel: rbf, linear | probability = True | 40 |
| **SVM_2** | C: 0.01, 0.1, 1, 10; class_weight: balanced, None; gamma: scale, 0.001, 0.01 | kernel = rbf; probability = True | 24 |
| **MLP_1** | alpha: 0.0001, 0.001, 0.01; hidden_layer_sizes: (50,), (100,), (50,25), (100,50); learning_rate_init: 0.001, 0.01; max_iter: 500, 1000 | activation = relu; early_stopping = True; solver = adam | 48 |
| **MLP_2** | alpha: 0.001, 0.01, 0.1, 1.0; batch_size: 32, 64, 128, auto; beta_1: 0.85, 0.9, 0.95; hidden_layer_sizes: (64,), (128,), (256,), (128,64), (256,128), (128,64,32), (256,128,64); learning_rate: constant, adaptive; learning_rate_init: 0.0001, 0.001, 0.01; max_iter: 500, 1000 | activation = relu; early_stopping = True; solver = adam | 4,032 |

**Table S6: Effect of MSA construction strategy on MSA composition and prediction performance of GEMME, PHACT, and PHACTboost.** Four MSA-generation configurations were evaluated combining the presence or absence of a BLAST-hit filtering step (Unfiltered/Filtered) and an MMseqs2-based clustering step (Full/Clustered). MSA size and identity to query are reported as the mean ± standard deviation across all proteins; identity to query was computed as the percentage of matching residues over all non-gap query positions. "Calculated" denotes the percentage of variants for which the respective tool returned a valid prediction; AUC values are computed only on successfully scored variants.

| Configuration | MSA size (mean ± std) | Identity to query (mean ± std) | GEMME | | PHACT | | PHACTboost | |
|---|---|---|---|---|---|---|---|---|
| | | | Calculated | AUC | Calculated | AUC | Calculated | AUC |
| **Full-Unfiltered** | 10227 ± 2881 | 44 ± 17 | 98.5% | 0.761 | 93.4% | 0.764 | 98.2% | 0.701 |
| **Full-Filtered** | 970 ± 152 | 71 ± 16 | 98.5% | 0.718 | 98.5% | 0.739 | 98.6% | 0.729 |
| **Clustered-Unfiltered** | 2680 ± 1396 | 40 ± 17 | 98.3% | 0.752 | 98.3% | 0.750 | 98.4% | 0.724 |
| **Clustered-Filtered** | 179 ± 124 | 70 ± 17 | 97.4% | 0.710 | 98.0% | 0.730 | 98.4% | 0.711 |

**Table S7: Overall performance of fourteen benchmarked VEPs on each tool's calculated entries from Benchmark dataset.** Tools are ranked by area under the receiver operating characteristic curve (AUC). "Datapoints" indicate the number of benchmark entries for which each tool successfully returned a prediction. Accuracy, precision, recall, F1 score, and Matthew's correlation coefficient (MCC) were computed using a classification threshold derived from Youden's J statistic, applied only to entries for which a given tool produced a valid prediction.

| Tool | Datapoints | AUC | Accuracy | Precision | Recall | F1 | MCC |
|---|---|---|---|---|---|---|---|
| VariPred | 10,384 | 0.773 | 0.703 | 0.766 | 0.625 | 0.689 | 0.418 |
| VESPA | 10,132 | 0.761 | 0.703 | 0.720 | 0.724 | 0.722 | 0.404 |
| GEMME | 10,210 | 0.752 | 0.696 | 0.712 | 0.719 | 0.716 | 0.390 |
| PHACT | 10,213 | 0.750 | 0.698 | 0.745 | 0.658 | 0.698 | 0.402 |
| MutPred2 | 10,384 | 0.748 | 0.691 | 0.721 | 0.671 | 0.695 | 0.383 |
| ESM1b | 10,384 | 0.739 | 0.688 | 0.751 | 0.606 | 0.671 | 0.388 |
| VESPAl | 10,328 | 0.738 | 0.686 | 0.706 | 0.692 | 0.699 | 0.370 |
| VespaG | 10,258 | 0.735 | 0.677 | 0.724 | 0.631 | 0.674 | 0.360 |
| PolyPhen-2 | 9,933 | 0.728 | 0.675 | 0.716 | 0.654 | 0.684 | 0.352 |
| PROVEAN | 10,385 | 0.725 | 0.684 | 0.672 | 0.777 | 0.721 | 0.365 |
| PHACTboost | 10,219 | 0.724 | 0.680 | 0.677 | 0.759 | 0.716 | 0.356 |
| EVE | 8,935 | 0.700 | 0.664 | 0.704 | 0.686 | 0.695 | 0.321 |
| PredictSNP | 10,385 | 0.683 | 0.687 | 0.680 | 0.761 | 0.718 | 0.371 |
| 3Cnet | 10,385 | 0.658 | 0.625 | 0.656 | 0.601 | 0.628 | 0.253 |

**Table S8: Stratified performance of the 14 benchmarked VEPs by data source and training-data independence level.** The evaluation was performed on subsets of each stratum, for which all 14 tools produced a valid prediction. N denotes number of datapoints in each stratum, for which all tools produced a prediction. AUC = area under the receiver operating characteristic curve. ClinVar and Humsavar represent clinically annotated human variants, whereas PMD and MMP represent functional, cross-species variant-effect data (see Methods). Independence level reflects each entry's relationship to the training data of the evaluated supervised VEPs (see Methods). Tools are listed in the order of Table 2.

| Tool | Full dataset | Data Source | | | | Independence Level | | |
|---|---|---|---|---|---|---|---|---|
| | | ClinVar | Humsavar | PMD | MMP | Clean sequence | Clean site | Clean variant |
| | N=8,622 | N=2,074 | N=1,829 | N=1,676 | N=3,043 | N=3,702 | N=4,486 | N=434 |
| | AUC | AUC | AUC | AUC | AUC | AUC | AUC | AUC |
| VariPred | 0.758 | 0.895 | 0.812 | 0.677 | 0.648 | 0.764 | 0.743 | 0.851 |
| VESPA | 0.755 | 0.853 | 0.765 | 0.722 | 0.705 | 0.772 | 0.743 | 0.773 |
| PHACT | 0.746 | 0.812 | 0.769 | 0.712 | 0.713 | 0.761 | 0.731 | 0.814 |
| GEMME | 0.737 | 0.828 | 0.759 | 0.707 | 0.693 | 0.737 | 0.734 | 0.797 |
| VESPAl | 0.734 | 0.783 | 0.739 | 0.718 | 0.711 | 0.746 | 0.729 | 0.754 |
| ESM1b | 0.732 | 0.860 | 0.764 | 0.642 | 0.660 | 0.723 | 0.734 | 0.771 |
| VespaG | 0.731 | 0.774 | 0.716 | 0.734 | 0.734 | 0.753 | 0.716 | 0.695 |
| MutPred2 | 0.729 | 0.842 | 0.799 | 0.633 | 0.637 | 0.699 | 0.742 | 0.801 |
| PolyPhen-2 | 0.722 | 0.794 | 0.738 | 0.697 | 0.670 | 0.704 | 0.737 | 0.744 |
| PHACTboost | 0.718 | 0.786 | 0.792 | 0.680 | 0.648 | 0.688 | 0.744 | 0.827 |
| PROVEAN | 0.708 | 0.821 | 0.728 | 0.665 | 0.670 | 0.692 | 0.723 | 0.709 |
| EVE | 0.700 | 0.761 | 0.684 | 0.666 | 0.701 | 0.705 | 0.694 | 0.709 |
| PredictSNP | 0.678 | 0.723 | 0.700 | 0.665 | 0.648 | 0.672 | 0.685 | 0.697 |
| 3Cnet | 0.631 | 0.798 | 0.732 | 0.498 | 0.526 | 0.640 | 0.602 | 0.777 |

**Table S9: Effect of excluding close training-set homologs on tool performance.** AUC of the 14 VEPs on the clean sequence stratum and on its homolog-free subset (see Supplementary Note 2), both restricted to the common subset. SD is the bootstrap standard deviation of the homolog-free AUC (1,000 stratified resamples). Tools are listed in the order of Table 2.

| **Tool** | **Clean sequence** | **Homolog-free** | |
|---|---|---|---|
| | N=3,702 | N=658 | |
| | **AUC** | **AUC** | **SD** |
| VariPred | 0.764 | 0.740 | 0.020 |
| VESPA | 0.772 | 0.749 | 0.019 |
| PHACT | 0.761 | 0.755 | 0.020 |
| GEMME | 0.737 | 0.741 | 0.020 |
| VESPAl | 0.746 | 0.724 | 0.020 |
| ESM1b | 0.723 | 0.653 | 0.022 |
| VespaG | 0.753 | 0.692 | 0.021 |
| MutPred2 | 0.699 | 0.684 | 0.021 |
| PolyPhen-2 | 0.704 | 0.666 | 0.022 |
| PHACTboost | 0.688 | 0.689 | 0.021 |
| PROVEAN | 0.692 | 0.682 | 0.021 |
| EVE | 0.705 | 0.665 | 0.021 |
| PredictSNP | 0.672 | 0.641 | 0.017 |
| 3Cnet | 0.640 | 0.576 | 0.023 |

**Table S2: Hyperparameter optimization of the final XGBoost classifier.** Hyperparameters were selected via 5-fold cross-validation grid search on the training set, optimizing validation AUC. The search space covered tree complexity (max_depth, min_child_weight, gamma), ensemble size (n_estimators), regularization strength (reg_alpha for L1, reg_lambda for L2), and stochastic subsampling (subsample for training rows, colsample_bytree for features per tree). XGB_2 denotes the initial grid configuration, while XGB_3 denotes the extended hyperparameter grid. Best value is highlighted in bold and reported in designated column.

| **Hyperparameter** | **XGB_2** | | **XGB_3** | |
|---|---|---|---|---|
| | **Grid** | **Best value** | **Grid** | **Best value** |
| colsample_bytree | [0.5, 0.6, **0.8**] | 0.8 | [0.8, 0.9, **1.0**] | 1.0 |
| gamma | [**0.1**, 0.5, 1.0] | 0.1 | [**0.05**, 0.1, 0.5] | 0.05 |
| learning_rate | [0.005, 0.01, **0.05**] | 0.05 | [**0.025,** 0.05, 0.075, 0.1] | 0.025 |
| max_depth | [2, 3, **4**] | 4 | [3, 4, **5**] | 5 |
| min_child_weight | [**5**, 10, 20] | 5 | [**3**, 4, 5] | 3 |
| n_estimators | [100, 200, **300**] | 300 | [300, 350, **400**] | 400 |
| reg_alpha | [**1.0**, 5.0, 10.0] | 1.0 | [**0.5**, 1.0, 1.5] | 0.5 |
| reg_lambda | [**2.0**, 5.0, 10.0] | 2.0 | [**2.0**, 5.0, 7.5, 10.0, 12.5] | 2.0 |
| subsample | [0.5, **0.6**, 0.8] | 0.6 | [**0.6**, 0.8, 0.9] | 0.6 |

**Table S3: Comparison of cross-validation performance of XGBoost classifiers trained using initial and extended hyperparameter grids.** AUC = area under the receiver operating characteristic curve. SD = standard deviation. CV = 5-fold cross-validation; CV — Train and CV — Val. report mean AUC and SD across folds on the training and held-out validation fold, respectively. XGB_2 denotes the initial grid configuration, while XGB_3 denotes the extended hyperparameter grid. Best value is highlighted in bold and reported in designated column.

| Model | CV - Train | | CV - Val. | |
|---|---|---|---|---|
| | AUC | SD | AUC | SD |
| **XGB_2** | 0.890 | 0.002 | 0.826 | 0.009 |
| **XGB_3** | 0.906 | 0.002 | 0.829 | 0.010 |

**Table S4: TreeSHAP and permutation feature importance (PFI) rankings for all nine constituent tools of the PredictSNPprot consensus model, on the training and test sets.** For each tool, rank, mean, and standard deviation (SD) are reported for both the mean absolute TreeSHAP value (log-odds contribution) and the mean permutation feature importance (mean ROC-AUC drop upon feature shuffling), evaluated independently on the preprocessed training and test partitions (see Methods). Tools are ordered by training-set TreeSHAP rank.

| Tool | Training set | | | | | | Test set | | | | | |
|---|---|---|---|---|---|---|---|---|---|---|---|---|
| | treeSHAP | | | PFI | | | treeSHAP | | | PFI | | |
| | Rank | Mean | SD | Rank | Mean | SD | Rank | Mean | SD | Rank | Mean | SD |
| **VariPred** | 1 | 0.581 | 0.004 | 1 | 0.067 | 0.003 | 1 | 0.584 | 0.008 | 1 | 0.057 | 0.006 |
| **PHACT** | 2 | 0.436 | 0.002 | 2 | 0.044 | 0.002 | 2 | 0.433 | 0.004 | 2 | 0.036 | 0.004 |
| **3Cnet** | 3 | 0.292 | 0.002 | 3 | 0.036 | 0.002 | 3 | 0.290 | 0.005 | 3 | 0.021 | 0.003 |
| **VespaG** | 4 | 0.222 | 0.002 | 6 | 0.022 | 0.001 | 4 | 0.223 | 0.004 | 6 | 0.011 | 0.002 |
| **VESPAl** | 5 | 0.183 | 0.002 | 5 | 0.023 | 0.001 | 6 | 0.174 | 0.004 | 5 | 0.014 | 0.002 |
| **ESM1b** | 6 | 0.174 | 0.001 | 4 | 0.024 | 0.001 | 5 | 0.176 | 0.002 | 4 | 0.015 | 0.002 |
| **PROVEAN** | 7 | 0.169 | 0.001 | 7 | 0.018 | 0.001 | 7 | 0.171 | 0.003 | 7 | 0.010 | 0.002 |
| **GEMME** | 8 | 0.129 | 0.002 | 8 | 0.017 | 0.001 | 8 | 0.126 | 0.003 | 8 | 0.006 | 0.002 |
| **PHACTboost** | 9 | 0.078 | 0.001 | 9 | 0.012 | 0.001 | 9 | 0.078 | 0.002 | 9 | 0.005 | 0.001 |
